# A first look at the genome structure of hexaploid ‘Black Mitcham’ peppermint (Mentha piperita L.)

**DOI:** 10.1101/2024.03.15.585188

**Authors:** Samuel Talbot, Iovanna Pandelova, B. Markus Lange, Kelly Vining

## Abstract

Peppermint, *Mentha xpiperita* L., is a hexaploid (2n = 6x = 72) and the predominant cultivar of commercial mint oil production in the US. This cultivar is threatened because of high susceptibility to the fungal disease Verticillium wilt, caused by *Verticillium dahliae*. This report details the first draft polyploid chromosome-level genome assembly for this mint species. The ‘Black Mitcham’ genome resource will broaden comparative studies of disease resistance, essential oil biosynthesis, and hybridization events within the genus *Mentha*. It will also be a valuable contribution to the body of phylogenetic studies involving *Mentha* and other genera that contain species with varying ploidy levels.

## Introduction

Mints (*Mentha* spp.) are commercially grown for their distilled monoterpene-rich essential oils. Extracts are used for aroma and flavoring in a wide variety of consumer products ranging from cosmetics to candies, medicine and dental care. The major species of import to industry are spearmints *M. spicata* (Native spearmint), *M*.*×gracilis* (Scotch spearmint) and peppermint (*M*. ×*piperita*). The peppermint cultivar ‘Black Mitcham’ has been the most widely-grown mint for much of the history of commercial production in the United States. First grown in England in the 1750s, then in colonial New England (Landing 1969), ‘Black Mitcham’ is highly prized for its oil quality and yield. However, this clonally-propagated herbaceous perennial is highly susceptible to the vascular wilt fungus *Verticillium dahliae*. ‘Black Mitcham’ is also male-sterile and not amenable to traditional crossing, meaning introgression of resistance from other mints is not an option. Irradiation of ‘Black Mitcham’ resulted in the release of two cultivars in the 1970s that showed higher field resistance to Verticillium wilt, Todd’s Mitcham and Murray Mitcham (Murray and Todd 1972; Todd *et al*. 1977), but these cultivars are not grown widely, and ‘Black Micham’ continues to dominate the acreage for a crop with peppermint-like oil characteristics.

*M*. ×*piperita* has a complex allohexaploid genome. It was first determined to be a natural hybrid of *M. aquatica* and *M. spicata* based on karyotype data (Harley and Brighton 1977), where chromosome numbers of *M*. ×*piperita* were either 2n=6x=66 or 2n=6x=72. Molecular marker studies have indicated that *M. spicata* is itself a hybrid of *M. longifolia* and *M. suaveolens* (Gobert *et al*. 2002, 2006). ‘Black Mitcham’s chromosome composition is 2n=6x=72.

With advances in long-read DNA sequencing technology, plus concurrent advances in chromatin-based methods for chromosome scaffolding, larger, more complex plant genomes are being assembled to whole-chromosome level, often including telomeres. This has been done for the accession of the diploid mint species used for genetic studies, *Mentha longifolia* CMEN 585 (PI 557767) (Vining *et al*. 2022). In that genome assembly, genes encoding key enzymes in the monoterpene biosynthesis pathway have been located on chromosomes. Still, polyploidy remains a challenge for chromosome scaffolding and haplotype phasing of plant genomes.

As a sterile allohexaploid, the genome of ‘Black Mitcham’ is presumed to have not undergone meiotic recombination since it was first cultivated. Therefore, its subgenome structure is expected to be preserved. Here, we present the first draft genome assembly and gene annotation of ‘Black Mitcham’ peppermint. It was produced through a combination of Pacific Biosciences HiFi sequencing and Omni-C-based scaffolding, and contains partially-resolved haplotypes.

## Materials and Methods

### Plant material

Clonally-propagated plant material from ‘Black Mitcham’ peppermint (PI 557937, local ID CMEN 133) was obtained from the USDA National Clonal Germplasm Repository in Corvallis, Oregon, USA. It was maintained at the Oregon State University West greenhouse complex by regularly transplanting a small wedge of soil containing stem and root material into fresh potting soil.

### Genome sequencing and assembly

High molecular weight genomic DNA was isolated from young, unexpanded leaf tissue using a modified cetyl trimethyl ammonium bromide method (Healey *et al*. 2014). Library preparation and sequencing on a Pacific Biosciences Sequel II instrument were done at Oregon CBD (Independence, OR). For high-throughput chromosome conformation capture (Hi-C), young, unexpanded leaf tissue was sent to Cantata Bio (Santa Cruz, CA, USA) for tissue processing, chromatin isolation and Omni-C library preparation. Libraries were sequenced as 150bp PE reads on an Illumina HiSeqX platform.

An initial genome size was estimated with a *K-mer* analysis of raw HiFi reads using jellyfish (version 2.3.0, RRID: SCR_005491) and the web version of GenomeScope (version 2.0, RRID: SCR_017014) with settings: *k-mer* length of 31, read length of 15,000 bp, and ploidy of six. Raw PacBio HiFi reads were assembled using hifiasm (version 0.16.1-r375, RRID: SCR_0121069) (Cheng et al., 2021). ‘Black Mitcham’ was assembled using the --primary flag. The Arima HiC mapping pipeline (https://github.com/Armia/Genomics/mapping_pipeline) was followed to map Omni-C reads to the respective genome assembly. To scaffold the contig assemblies into pseudo-chromosomal scaffolds, YaHs (version 1.1, RRID: SCR_021172) was run using the Hi-C aligned, read name sorted bam file (Zhao et al., 2022). A Hi-C contact map was generated and manually curated by contig merging and rearrangement within JuiceBox (version 1.11.08). This was done to obtain an accurate number of pseudomolecules for the species based on expected pseudomolecule size according to alignments of *M. longifolia* and suggested protocols by Aiden et al. (2021). Final assembly metrics were generated by a custom script; assembly completeness was assessed with BUSCO (version 5.5.2, RRID: SCR_015008) in genome mode, using the Embryophyta_odb10 dataset (Manni et al., 2021). To assess long-range structural variations, ‘Black Mitcham’ was compared to the collapsed diploid assembly of *M. longifolia* V3 using minimap2 with divergence rate set to 20 percent (version 2.23-r1111, RRID: SCR_018550), SyRI (version 1.6.3, RRID: SCR_023008) and visualized by plotsR (Goel *et al*. 2019; Li *et al*. 2021; Goel and Schneeberger 2022).

### Repeat identification

To create a repeat library of transposable element families, EDTA (version 2.1, RRID: SCR_XYZ) was used on the finalized genome assembly with parameters: --anno 1 --sensitive (Ou et al., 2019). EDTA is composed of a number of tools, including TIR-Learner, Generic Repeat Finder, HelitronScanner and TEsorter. Low complexity DNA sequences and repetitive regions identified by EDTA were soft masked prior to gene annotation. The quality of the genomes assembled repetitive regions were assessed using the long terminal repeat (LTR) assembly index (LAI); this pipeline was composed of LTRharvest in GenomeTools (version 1.6.1, RRID: SCR_016120), LTR_FINDER (version 1.2, RRID: SCR_015247), and LTR_retriever (version 2.9.6, RRID: SCR_017623) using suggested parameters to predict and combine likely full length candidate LTR-RTs (retrotransposons) (Gremme et al., 2013; Ellinghaus et al., 2008; Xu and Wang, 2007; Ou and Jiang 2019; Su et al., 2019; Shi and Liang, 2019; Xiong et al., 2014; Zhang et al.., 2022; Ou et al., 2018). Calculation of the LAI index was based on the formula:

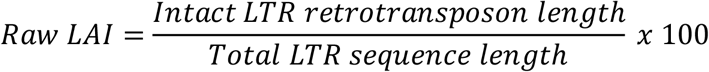

### Structural gene annotation

Gene prediction and annotation was facilitated by 150bp PE Illumina RNA-Seq data derived from ‘Black Mitcham’ root and stem tissue (unpublished), and concatenated with glandular trichome from *M. xpiperita* (Ahkami et al., 2015) and flower tissue (Boachon et al., 2018) for a total of ∼380 million reads. Structural annotation of protein-coding genes were identified using the gene prediction software AUGUSTUS and GeneMark-ETP+, integrated by BRAKER3 (Stanke et al., 2006, 2008; Gotoh 2008; Buchfink et al., 2015; Hoff et al., 2016, 2019; Kovaka et al., 2019; Pertea and Pertea 2020; Brůna et al., 2021, 2023). First, RNA-Seq reads were aligned to the chromosome-level assembly using the splice-aware aligner Hisat2 (Kim et al., 2019) to generate a bam file. Second, BRAKER3 was run using the soft masked genome output from EDTA, the RNA-Seq bam file, and a curated Viridiplantae odb11 protein set. For homology-based annotation, a protein set derived from the previous *M. longifolia* genome assembly (Vining et al., 2022) and two unpublished Mentha diploid species assemblies consisted of 187,098 proteins, and was used as input to BRAKER2 with GenomeThreader (Gremme 2013) with AUGUSTUS ab-initio and GALBA (Brůna et al., 2023; Li 2023), separately. Gene predictions of the respective BRAKER3 and homology-based annotation runs were assessed for quality, deduplicated, and combined using TSEBRA with default settings (-c braker3.config) (Gabriel et al., 2021). Completeness of the predicted annotation sets were assessed using BUSCO protein mode with the Embryophyta odb10 data set. The proteome set was additionally vetted with OMArk (version 0.3.0) using default settings, the OMAmer Viridipla ntae v2.0.0.h5 database, and a .splice file representing all alternatively spliced variants of transcripts (Nevers et al., 2024).

### Functional gene annotation

A single “best” haplotype was selected for functional analysis, consisting of 12 pseudo-chromosomal scaffolds with the highest quality alignment to *M. longifolia* V3. This set of chromosomes and their respective annotated gene models were subject to predictive functional analysis using within OmicsBox (version 3.1); the OmicsBox pipeline included CloudBLAST using BLASTx, InterPro, GO Merge, GO Mapping, and GO Annotation plus validation (Götz et al., 2008; Paysan-Lafosse et al., 2023). Conservation of putative high confidence gene models between assemblies was assessed using OrthoFinder (version 2.5.4, RRID: SCR_017118) (Emms and Kelly 2019).

## Results and Discussion

### Genome assembly

A total of 1.9 million PacBio HiFi reads with an average length of 18,304 bp were generated from one 8M SMRT cell, resulting in 34.8 Gb of sequencing data for *M*. ×piperita ‘Black Mitcham’ (Supplement Table S1). *K-mer* counts of the hexaploid ‘Black Mitcham’ identified three peaks and a haplotype genome size estimate of 342 Mb (Supplement Figure S1). ‘Black Mitcham’ was assembled using hifiasm and the --primary flag, generating 610 contigs for a total of 2,040 Mb and an N50 of 27.7 Mb (Table 1), representing 84% coverage of the 2.4 Gb expected genome size.

**Table 1.** Summary statistics for the *M. longifolia* and ‘Black Mitcham’ peppermint genome assemblies.

| Metrics | MlongV3 | BlackMitcham |
| --- | --- | --- |
| Size (Mb) | 469 | 2,040 |
| Number of contigs | 37.5 | 610 |
| Size (chr only, Mb) | 462.6 | 1,974 |
| N50 (Mb) | 37.5 | 27.7 |
| Longest length chr (Mb) | 46.6 | 42 |
| BUSCO (genome) | 97.8% | 99.5% |
| LAI score | 17.9 | 10.1 |
| Number of Protein Coding Genes | 42,107 | 247,493 |

The hifiasm haplotype assemblies were inputs to the chromosome scaffolding process. The Omni-C protocol (Cantata Biosciences) was used to generate the sequencing library which was then scaffolded using YaHs (version 1.1), and manually curated within JuiceBox. Approximately 40 million PE 150 bp reads were generated for Hi-C, for a total yield of ∼12 Gb (6x coverage) (Table S1). The scaffolded chromosome-level assembly spans 1,974 Mb, capturing 96% of the hifiasm primary assembly. Following manual-curation within JuiceBox based on minimap2 alignments to *M. longifolia* V3, 72 pseudo-chromosomal scaffolds were generated. BUSCO results in genome mode showed that the assembly was high quality and captured >98% of conserved genes in the embryophyta odb10 dataset (Table 1).

### Characterization of repeats

To further vet the genome assembly, the LTR Assembly Index was calculated, wherein a LAI score less than 10 indicates draft quality, 10-20 as reference quality, and gold standard with an LAI score greater than 20 (Ou et al., 2022). Subsequent LAI analysis of the reference *M. longifolia* V3 was estimated at 17.9, whereas the LAI score of ‘Black Mitcham’ was 10.13 and indicative of a draft genome assembly (Table 1).

Prior to gene annotation, genomic repeats were identified on all assemblies for downstream masking. The proportion of repeats and unknown elements identified by EDTA resulted in ‘Black Mitcham’ having ∼62% of its genome masked (Supplement Table S2). This proportion of repeats in ‘Black Mitcham’ is significantly higher than *M. longifolia* V3 which had ∼45% of the genome masked. The majority of repeats were Class I Long Terminal Repeats (LTRs), with 32.3% of the genome composed of the Copia or Gypsy superfamily. A significant proportion of the genome (12.5%) consisted of helitron elements. Helitrons are unique transposable elements that are thought to significantly contribute to genome rearrangements, gene fragment duplications, exon shuffling and chimeric transcript generation (Barro-Trastoy and Köhler, 2024). Published helitron content estimates vary widely among plant species and between genotypes within a species; in the hexaploid wheat cultivar ‘AK58’, helitrons made up 6% of total genome size whereas in ‘Chinese Spring’ they made up only 0.047% (Wang *et al*. 2022). Such a difference in helitron density could be attributed to computational difficulty in distinguishing helitron elements that are embedded in other repetitive sequences.

### Genome annotation

For ‘Black Mitcham’, 247,493 protein-coding genes were annotated in the genome assembly, representing roughly ∼41,200 genes per haplotype. The number of gene models in ‘Black Mitcham’, when considering an individual haplotype, is comparable to the unphased *M. longifolia* V3 diploid assembly annotation of 42,107 genes. Translated coding sequences of putative gene models were subject to BUSCO analysis and revealed that the annotation set was 99.4% complete, although nearly all complete copies were duplications, which is expected due to polyploidy (Table 2). As an additional check, OMARK was used to compare *M. longifolia* V3 to ‘Black Mitcham’. The analysis showed that the majority of conserved hierarchical orthologous groups derived from the lamiids clade were duplicated in *M. x piperita*, with ∼36% of proteins having unknown lineage placement (Figure 1). A set of twelve Black Mitcham scaffolds with the best alignment to the twelve *M. longifolia* V3 chromosomes was chosen to represent one Black Mitcham haplotype. This haplotype contained 43,758 gene models, which were then used as input to OmicsBox for functional annotation. A total of 32,765 (∼75%) gene models was given a functional description (Supplement Figure 2).

**Table 2.** Assessment of annotation completeness in *M. longifolia* and ‘Black Mitcham’.

| Genome | Protein categories | BUSCO |  |
| --- | --- | --- | --- |
|  |  | Number | Percentage |
| *M. longifolia* (V3) | Complete BUSCOs (C) | 1413 | 87.5 |
|  | Complete and single-copy BUSCOs (S) | 1140 | 70.6 |
|  | Complete and duplicated BUSCOs (D) | 273 | 16.9 |
|  | Fragmented BUSCOs (F) | 95 | 5.9 |
|  | Missing BUSCOs (M) | 106 | 6.6 |
|  | Total BUSCO groups searched | 1614 | 100.0 |
| Black Mitcham (primary) | Complete BUSCOs (C) | 1605 | 99.4 |
|  | Complete and single-copy BUSCOs (S) | 10 | 0.6 |
|  | Complete and duplicated BUSCOs (D) | 1595 | 98.8 |
|  | Fragmented BUSCOs (F) | 3 | 0.2 |
|  | Missing BUSCOs (M) | 6 | 0.4 |
|  | Total BUSCO groups searched | 1614 | 100.0 |

**Figure 1.**
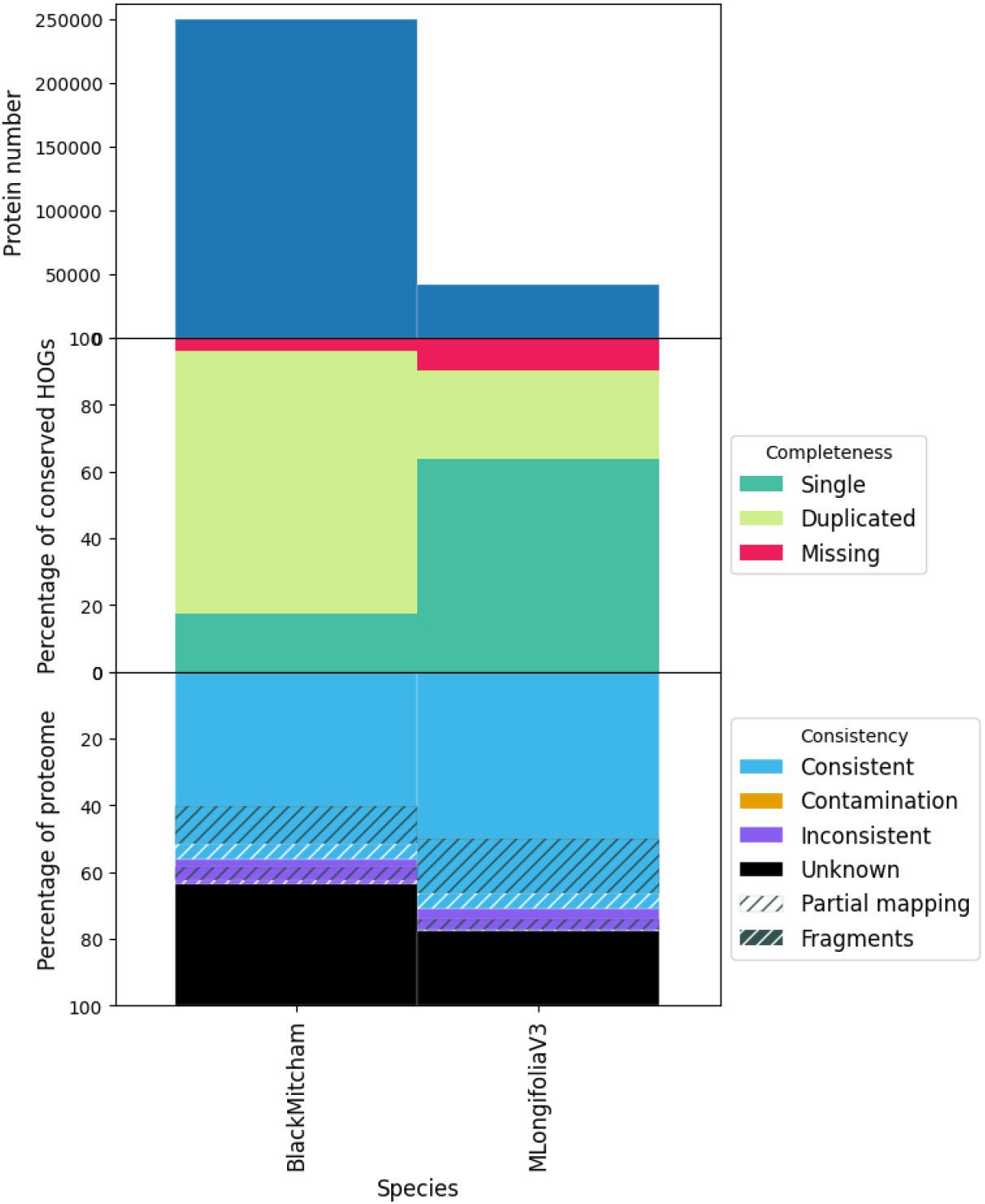
OMArk results of Gene annotation sets of *M. longifolia* and ‘Black Mitcham’.

**Figure 2.**
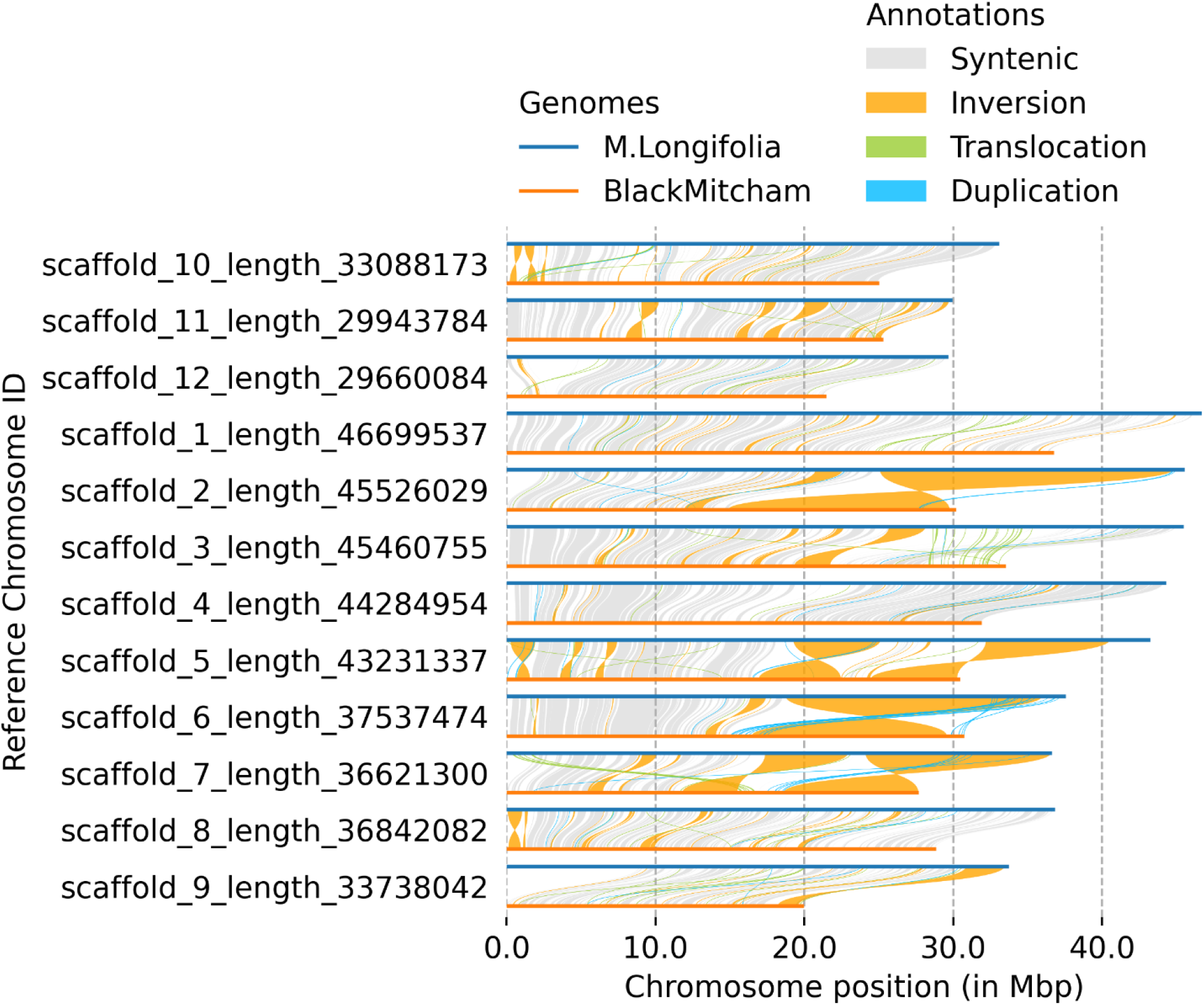
SyRI plot of twelve ‘Black Mitcham’ pseudo-chromosomes against the *M. longifolia* V3 genome assembly.

To assess gene models with shared homology, OrthoFinder (version 2.5.4, RRID: ) was run with the annotated amino acid sequences derived from *M. longifolia* V3 and the ‘Black Mitcham’ gene set. Approximately 77% (223,735) gene models were assigned orthogroups were categorized into orthogroups, with ∼63% of these being orthogroups with both species present (Supplement Table S3). Of the species-specific orthogroups, the large majority (67%) of these orthogroups were composed of a single gene model. Nearly 66,000 (23%) gene models were unassigned. The large number of single copy orthogroups and unassigned gene models is indicative of both novel gene models and highly fragmented genes.

### Synteny between ‘Black Mitcham’ and M. longifolia

As an initial step to identify syntenous regions between ‘Black Mitcham’ and its hypothesized ancestral species, the twelve highest-quality ‘Black Mitcham’ pseudo-chromosome scaffolds were aligned to those of *M. longifolia* V3 using minimap2 and SyRI, then pairwise alignments were plotted using plotsr. Only ∼40% of the total length of the ‘Black Mitcham’ scaffolds aligned meaningfully to those of *M. longifolia* (Supplement Table S4, S5). Of the syntenous regions between the two genomes, the majority of alignments contained inversions (Figure 2). These results were expected for several reasons. First, the ‘Black Mitcham’ subgenome haplotypes are only partially resolved, and are not defined in any way based on hypothesized ancestry. Second, the specific representatives of the hypothesized ancestral species subgenome contributors are unknown. Third, the *M. longifolia* diploid ‘CMEN 585’ is from South Africa and is assumed to not be a direct ancestor of ‘Black Mitcham’ which reportedly arose in Europe.

The results reported here open the door to further exploration of the origin of ‘Black Mitcham’ peppermint, including definition of subgenomes by species-of-origin, as has been done for the allo-octoploid cultivated strawberry, *Fragaria* ×*annanassa* (Edger *et al*. 2019). However, unlike the sexually fertile cultivated strawberry, the chromosomes in the subgenomes of ‘Black Mitcham’ are presumed to be frozen in time, awaiting discovery.

### Data Availability

The primary assembly and related annotation will be available through mintgenomes.oregonstate.edu as a fully accessible genome browser (in progress). These files with additional scripts that were used to assemble the genome are also accessible through our Github at https://github.com/ViningLab/ViningLab_code/blob/3289b81866d0e01c4fa824e460c42e2f0c5b126e/G3_BM_AssemblyPipeline.

## Acknowle dge ments

We thank Cantata Biosciences for their collaboration on genome scaffolding for this project. This work was partially funded by the Mint Industry Research Council.

